# Cerebral Topographies of Perceived and Felt Emotions

**DOI:** 10.1101/2023.02.08.521183

**Authors:** H Saarimäki, L Nummenmaa, S Volynets, S Santavirta, A Aksiuto, M Sams, IP Jääskeläinen, JM Lahnakoski

## Abstract

Emotions modulate behavioral priorities based on exteroceptive and interoceptive inputs, and the related central and peripheral changes may often be experienced subjectively. Yet, it remains unresolved whether the perceptual and subjectively felt components of the emotion processes rely on shared brain mechanisms. We applied functional magnetic resonance imaging, a rich set of emotional movies, and high-dimensional, continuous ratings of perceived and felt emotions depicted in the same movies to investigate their cerebral organization. Eight main dimensions of emotions evoked during natural movie scene perception were represented in the brain across numerous spatial scales and patterns. Perceived and felt emotions generalized both between individuals and between different samples of stimuli depicting the same emotions. The neural affective space demonstrated an anatomical gradient from responses independent of specific emotions in polysensory areas and default mode regions to more localized and emotion-specific discrete processing in subcortical regions. Differences in neural activations during felt and perceived emotions suggest that temporoparietal areas and precuneus have a key role in computing the affective value of the sensory input. This affective value is then transformed into the subjective emotional experience in the anterior prefrontal cortices, cerebellum, and thalamus. Altogether these data reveal the similarities and differences of domain-general and emotion-specific affect networks in the brain during a wide range of perceived and felt emotions.

## Introduction

Emotions promote survival by monitoring external and internal challenges and coordinating automatic changes in peripheral physiology, behavior, motivation, and conscious experiences (i.e., feelings) ^1,2,3^. Emotion circuits span the whole brain and are focused in the limbic and paralimbic structures ^4^. Despite significant advances in mapping the neural basis of emotions, two critical questions remain unanswered. First, how distinct emotional states - such as anger, fear, or disgust - are coordinated in the brain, and second, what is the relationship between the brain circuits that extract and recognize emotional information from the external input and those that subsequently generate the phenomenological experience of emotions.

Emotions are organized categorically across multiple levels ranging from brain activity ^5–9^, autonomic activation ^10^, somatosensory and interoceptive experience ^11^, facial, bodily, or vocal behaviors ^12–14^ to subjective feelings ^15^. Recent behavioral studies suggest that the human affective states span almost 30 distinct emotions ^16^. Yet, most imaging studies have investigated either lower-order emotional dimensions (valence and arousal) ^17^ or the canonical six “basic” emotions ^5,18–21^, only rarely extending the emotion space beyond these categories^6,22,23^.

Emotional processing begins with the encoding of the survival impact of sensory information based on, for example, the presence of predators or virus vectors, or others’ emotional expressions and behavior ^13,24,25^ and contextual cues^26^. These cascading evaluative processes may trigger the central and peripheral emotional responses and subsequently also the subjective experience or feeling of emotion (see, e.g., ^1,3,27^). Emotional perception and feelings are associated with activation in overlapping brain regions ^28–30^. However, an event may trigger emotional states that are either congruent (e.g., seeing a smiling baby makes you happy) or incongruent (e.g., seeing a smiling villain makes you scared) with the event. Similarly, in anxiety disorders, emotional perception might be intact while the resulting feeling is disproportionally biased towards anxious or fearful feelings ^31^. Thus, to map the neural basis of emotional perception and feelings, we need to establish accurate stimulus models distinguishing these two aspects independently. Yet, human neuroimaging studies rarely differentiate perceptual versus experiential emotion processes ^32^.

Movies provide a powerful way to portray a wide range of human emotions. The characters may be involved in nuanced and powerful emotional states (such as Rick Blaine on the airstrip at the end of *Casablanca* or Rose DeWitt Bukater onboard the sinking passenger liner in *Titanic)*, and such scenes also elicit strong and consistent emotions in the viewers ^33–35,36^. Thus, carefully curated cinematic stimuli allow mapping the high-dimensional representation of both perceived and experienced emotions in the viewer’s brain.

In this study, we modeled the high-dimensional organization of perceived and felt emotions in the human brain and tested how these models generalize across stimuli and participants. We showed our participants two hours of emotional movies while measuring their brain activity with functional magnetic resonance imaging. We selected a wide range of emotion categories to cover a high-dimensional affective space (**Figure 1A**) and 2 hours of emotional movie scenes from an existing emotion-elicitation database (**Figure 1B**; ^37^). We fitted emotion feature models derived from the dynamic emotion ratings to the fMRI data and adopted a cross-validation scheme to evaluate the generalizability of the emotion models across participants and movie stimuli (**Figure 1C-D**). This allowed us to directly compare the neural basis of perceived and felt emotions using the same stimuli. The results show that movies elicited consistent neural responses to a wide range of perceived and felt emotions that generalize across stimuli and participants. The cerebral topographies of perceived and felt emotions were partly separate, and neural emotion clusters only partially correspond to behavioral emotion clusters.

**Figure 1.**
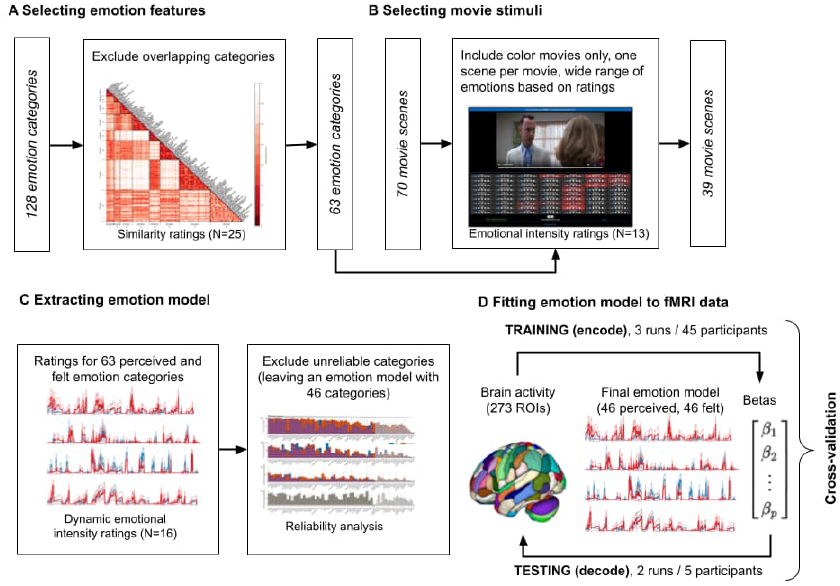
Methodological pipeline. **A**. Candidate emotions were selected based on previous studies. **B**. A total of 70 movie clips with total duration of >2 hours were selected from ^37^ to elicit a wide range of emotions. **C**. Dynamic ratings for perceived and felt emotion features were collected during movie viewing. The reliability of the ratings of the emotion features was evaluated and only the 46 most reliable features were included in the stimulus models. **D**. The emotion models were fit to the BOLD data using a cross-validation scheme.

## Results

### Reliability of ratings of perceived and felt emotions

Subjective ratings confirmed that the participants perceived and felt a wide array of different emotions while viewing the movies (**Figure 2A-C**). However, there was significant variation in both occurrence, intensity, and intersubject consistency of the perceived and felt emotions. We found a clear continuum from commonly and consistently reported emotions (*fear, anxiety, despair, devastated, sadness*) to infrequently and inconsistently reported emotions (*jealousy, craving, hurt, satisfaction*). The mean intersubject correlation of the ratings and the number of participants giving above-zero ratings for each emotion were positively correlated. To avoid spurious effects, we show the clustering analysis (**Figure 3**) and all further analyses only for the 46 emotions, which showed reliable, non-zero rating profiles across the five runs (**Figure 2**). Clustering analysis for all emotions further confirmed that unreliable emotions are separated from reliable ones (**Supplementary Figure S2**). Clustering for this reliable set of emotions revealed a broad valence-related structure. The nine clusters were formed around the following emotion categories: 1) sadness, anger, and fear, 2) disgust, 3) displeasure, 4) embarrassment, 5) love, 6) surprise, 7) gratitude and joy, 8) amusement, and 9) calmness.

**Figure 2:**
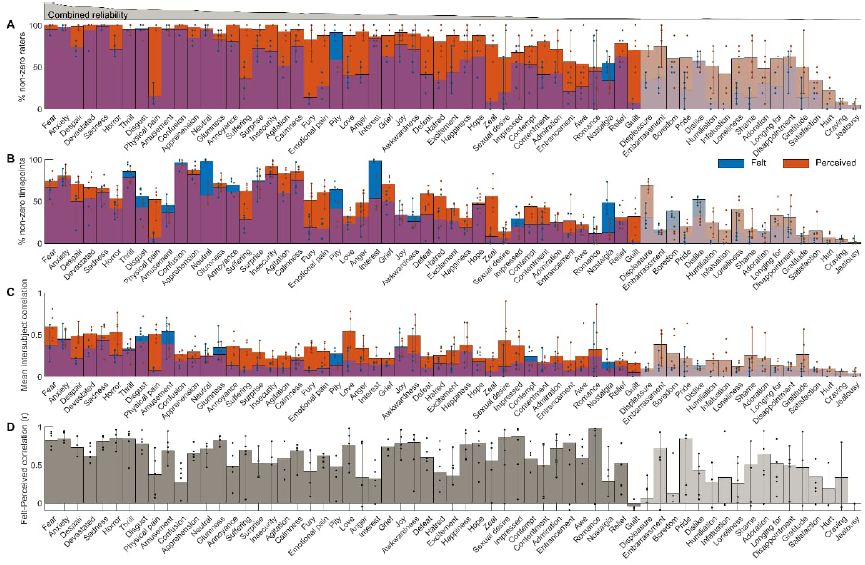
Reliability of emotion ratings. **A**. Percentage of raters for each emotion and each task (perceived and felt combined) with above-zero ratings for at least one time point. **B**. Percentage of time points for each emotion where at least one rater gave a non-zero rating. **C**. Mean intersubject correlation of emotion ratings across pairs of raters. **D**. Similarity of mean ratings for perceived and felt emotions across raters across all runs. In each panel, bars indicate the mean values across runs, dots show the values for individual runs, and vertical lines indicate the min-max range. The emotions are ordered based on the combined reliability of panels A–C calculated as the geometric mean of the percentage of non-zero raters, non-zeros timepoints, and intersubject correlation or ratings (shown in the gray plot at the top). Emotions not deemed reliable based on permutation statistics are shown with transparent colors.

**Figure 3.**
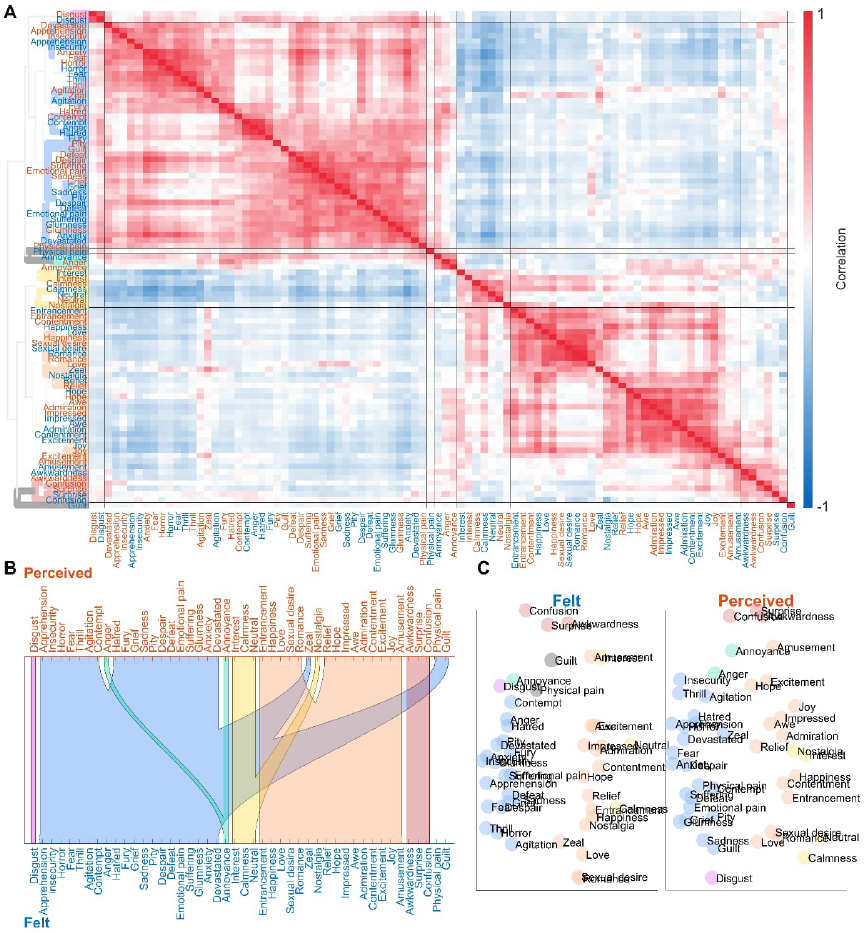
The cluster structure of perceived and felt emotions. **A:** Correlation matrix and dendrogram of ratings over emotion models across all runs and both felt and perceived emotions for the reliably occurring emotions. Colors of the dendrogram indicate the clustering solution based on rating data and is also reflected in subsequent figures. **B:** Alluvial diagram shows corresponding cluster structures between felt and perceived emotions. **C:** Multidimensional scaling visualizes the similarity of emotions based on the ratings of felt emotions (left) and perceived emotions (right). Emotions not belonging to any cluster are shown in gray in the dendrogram and multidimensional scaling plots and are left unconnected in the alluvial diagram.

### Temporal similarity between perceived and felt emotions

Next, we addressed the temporal similarities of perceived and felt emotions by calculating the similarity between the time series for the perceived and felt emotion from the same category (**Figure 2D**). Overall, the temporal similarity structures for the perceived and felt emotions were consistent. Out of the 63 emotions, the mean correlation between perceived and felt emotions was above .60 (corresponding to 36% shared variance) for 32 emotions categories. The emotions with the highest similarity included *romance, impression, sexual desire, pride*, fear-related emotions (including *fear, anxiety, horror*, and *thrill*), and sadness-related categories (including *sadness* and *glumness*). While the perception of emotion led to a consistent feeling of the same emotion for most emotion categories, there were also some emotions with discordant perception and feeling (e.g., *guilt, physical pain, confusion, anger, hatred, interest, excitement*, and *nostalgia*). The correspondence of the cluster structures between perceived and felt emotions further highlights this decoupling (alluvial diagram in **Figure 3**): Emotions clustering around *love, pride, sadness*, and *impression* had similar perception and feeling structures, whereas social emotions clustering around *humiliation, loneliness*, and *insecurity* were less consistently perceived and felt. While some emotions were perceived more often than felt (e.g., *physical* and *emotional pain, fury, zeal*, and *guilt*), others such as *pity* and *nostalgia* were mostly felt and rarely perceived in the movies (**Figure 2A**).

### Brain basis of perceived and felt emotions

We first evaluated where in the brain different perceived and felt emotions are represented consistently across stimuli and participants (**Figure 4**). First, each of the five runs in our experiment contained different movies depicting the same set of emotions, allowing us to test the generalizability of emotion-related brain responses across stimuli. **Figure 4A** shows the generalizability (i.e., correlations) of the cross-validated emotion models where three runs were used for training and two runs for testing (for region-wise generalizability, see **Supplementary Table S1**). For both perception and feeling, the responses were most consistent in the right temporal cortex, bilateral temporoparietal junction, anterior cingulate cortex, and precuneus. For emotional perception, the responses were also consistent bilaterally in the temporal cortices. For feelings, the model was also consistent in the anterior, medial, and dorsolateral prefrontal cortices, thalamus, and cerebellum. Second, we evaluated the consistency of emotion-related brain responses across participants. **Figure 4B** shows that we found largely overlapping, widespread effects for both perception and feeling in regions covering occipital, temporal, and parietal lobes, posterior midline, and cerebellum. The only notable difference was the more consistent prefrontal activity for feelings, which was absent for perception.

**Figure 4.**
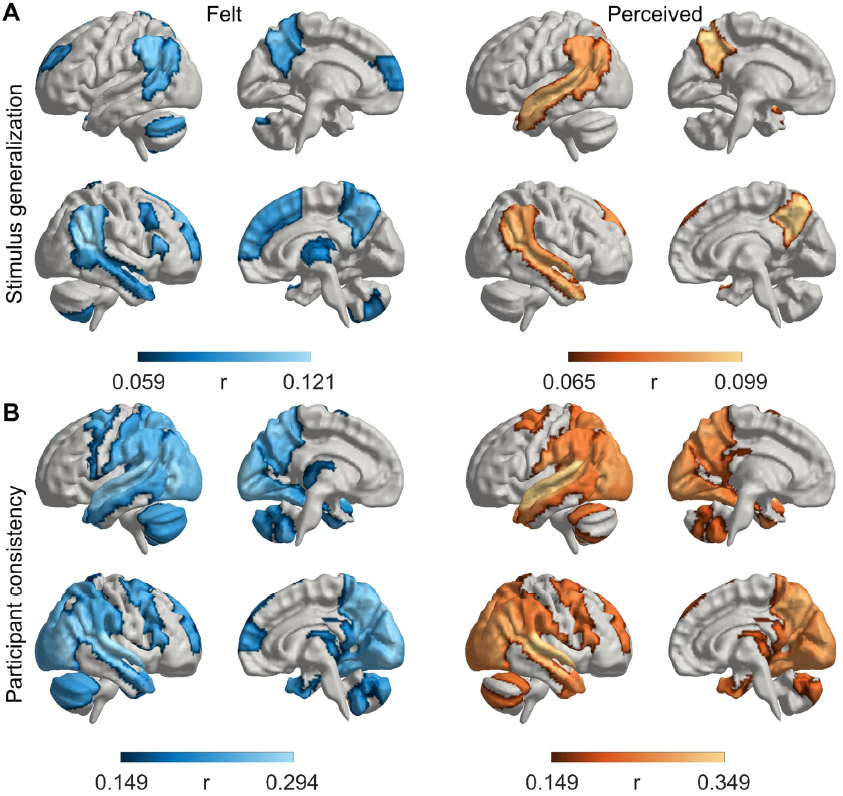
Consistency of emotion models. **A**. Generalizability of emotion responses for different stimuli for felt (left) and perceived (right) emotions. **B**. Consistency of the cross-validated models across participants for felt (left) and perceived (right) emotions. Regional data are thresholded at FWER p<.05 of the mean fit of all run combinations (3 training runs, 2 test runs, 10 combinations). Statistics are based on the 95th percentile of maximum statistics in surrogate data (max. over ROIs and 95th percentile over null iterations with circularly shifted training data).

To test whether low-level stimulus features could explain the emotion-related brain activity, we ran the same analysis with a model that included the emotion features and three low-level visual and auditory features (**Supplementary Figure S4**). As expected, low-level stimulus feature models explained sensory activity in V1/V2, V2/MT, and A1/STG. However, activity in regions, including TPJ, STS, and MPFC, was unaffected, showing that stimulus-related activity in these regions reflects emotional processing rather than purely sensory processing.

### Regional responses to specific perceived and felt emotions

Next, we analyzed ROI-level data to address the representation of perceived and felt emotions in different brain regions by quantifying the region-wise consistency of the responses evoked by each perceived and felt emotion (**Figure 5**). We first identified the emotion categories associated with the most widespread activity changes across the brain (**Figure 5**, bar plot). We found a clear gradient in the brevity of the distribution of emotion-evoked activity in the brain across emotions. Perceived and felt *amusement, thrill*, and *horror* led to the most widespread brain activation across the temporo-occipital, parietal, and midline regions. In contrast, more neutrally-valenced emotions, such as *calmness, neutral*, and *nostalgia*, yielded focal deactivation in parietal midline and lateral regions. Anger- and disgust-related emotions were also accompanied by deactivation in regions including STG and pSTS. Overall, sadness- and happiness-related, calm emotions yielded weaker responses in fewer areas. Feelings led to more widespread activation than perception for some emotions, such as *surprise, fear*, and *sexual desire*, while the opposite was true for other emotions, such as *agitation, zeal, annoyance, hatred, guilt, anger*, and *excitement*.

**Figure 5.**
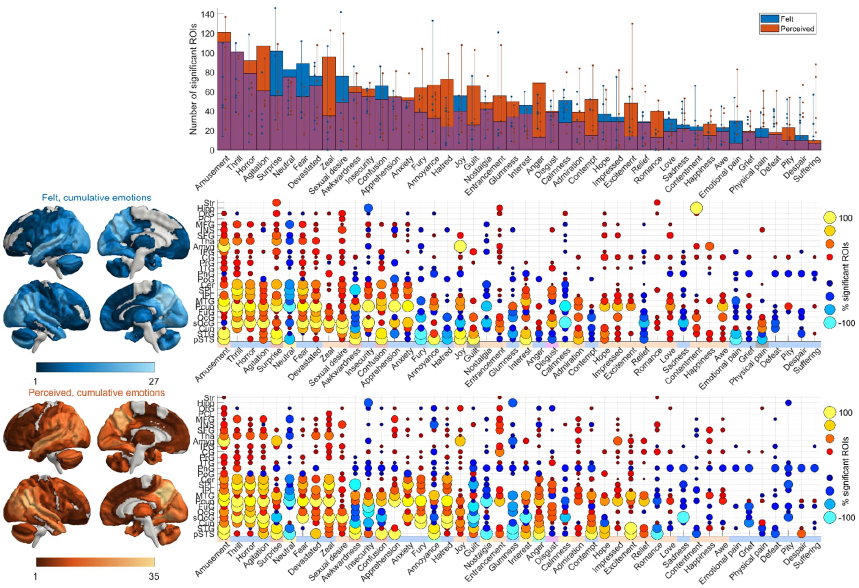
Regional responses to specific experienced and perceived emotions. Total brain area activated by individual emotions (**top**) and the regional consistency of emotion-wise responses for felt (**middle**) and perceived (**bottom**) emotions. Emotions are ordered based on the spatial extent of significant activations averaged over both tasks, and brain regions are ordered by the number of emotions that are significantly correlated with the regional activity. The bar plots show the average number of ROIs that were activated over all runs and the dots show individual-run results. Vertical lines connect the minimum and maximum values over runs. The brain plots show the number of emotions whose responses were statistically significant in at least one run; the matrix plot shows the corresponding data at the level of single emotions at the scale of macroanatomical regions, each comprising multiple ROIs across both hemispheres. The dots indicate the percentage of ROIs within the macroanatomical regions that were significantly activated (hot colors) or deactivated (cold colors) by the emotions over all the runs and over participants. The data are thresholded at two-tailed p < .05 (FWER corrected). For brain plots with higher resolution and bar plots color-coded with clustering results, see Supplementary Figure S5.

There were also apparent regional differences in the breadth of tuning for different emotions as demonstrated by the spatial gradient ranging from regions activated during most emotions to those activated during few specific emotions (**Figure 5**). Superior temporal and posterior midline regions (including precuneus and superior occipital gyrus) generally responded to most emotions for both perception and feeling. More selective activity to clusters of emotions was seen in regions such as cerebellum whose activity was associated with fear- and surprise-related emotions and in superior parietal regions which yielded consistent responses to fear-related emotions. Finally, regions such as the amygdala showed emotion-specific responses: in the amygdala activity was consistent especially for *amusement, joy*, and perceived *surprise*.

### Spatial clustering of the neural responses to felt and perceived emotions

We next analyzed the similarity of the response profiles by clustering the brain activity patterns based on their between-run similarity and compared it with the similarity of ratings (depicted in **Figure 3**). The clusters identified from the neural data differed from those in the self-report data (**Figure 6**, see **Supplementary Figure S3** for a side-by-side comparison of the similarity structures). The large valence-based (pleasure/displeasure) cluster structure in self-report data was absent in the neural data. At the neural level (**Figure 7**), the most prominent clusters contained amusement- and confusion-related states (cluster 4) and fear--related states (cluster 3). Felt *joy* drove a small cluster (cluster 6). Neutral and calm romantic emotions also formed another small cluster (cluster 7). Less reliable, but robust clusters were also found for *disgust* (cluster 2) and felt calm, positive emotions (cluster 8). A large set of emotions also stayed relatively independent of the larger clusters, forming two somewhat unreliable clusters, driven by perceived *happiness* and emotions with a clear bodily component, such as *physical pain* and *sexual desire* (clusters 5 and 1). Fear-related perceived and felt emotions consistently clustered together. Similar grouping was also apparent for positive and neutral emotions such as *amusement* and *confusion*.

**Figure 6.**
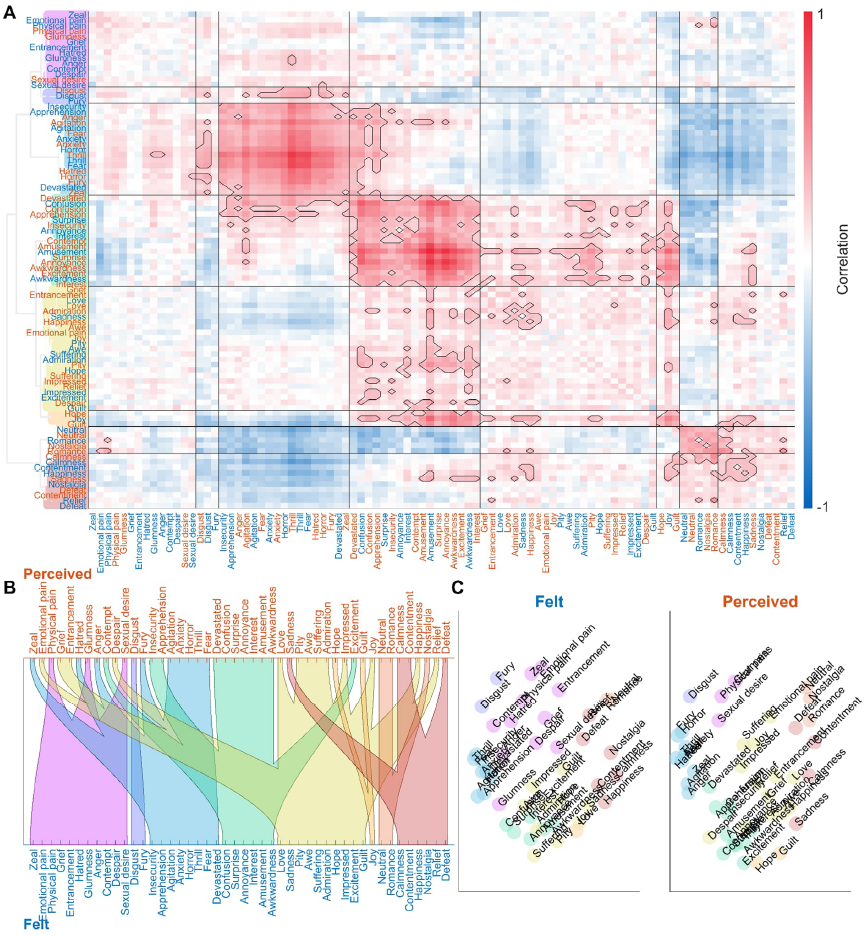
The cluster structure of the brain responses to perceived and felt emotions. The organization of the figure is similar to the clustering of ratings in Figure 3. **A:** Between-runs correlation matrix and dendrogram clustering of brain responses. **B:** Alluvial diagram of cluster correspondence between perceived and felt emotions. **C:** Multidimensional scaling of similarity of brain responses to felt (left) and perceived emotions (right).

**Figure 7.**
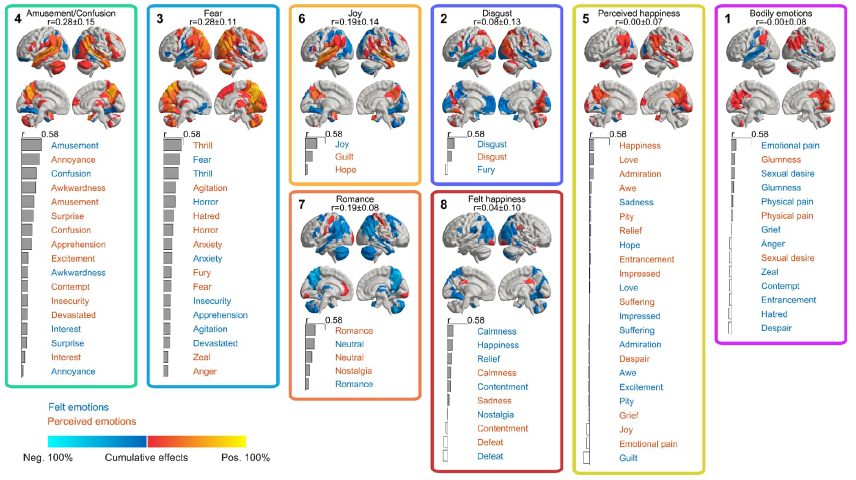
Brain-based clustering of emotions. Cumulative activity patterns for the brain clusters of perceived and felt emotions. Cluster colors and numbers depict the clustering hierarchy and order of emotion ratings in Figure 6. The clusters are ordered based on the mean reliability of the activity patterns (between-runs spatial correlation) of the emotions contained within the cluster. The emotions in the clusters are ordered by reliability, which is depicted by the bar plots. Brain maps show the cumulative significant (p<.05 Bonferroni corrected over regions and emotions) activations over runs and emotions belonging to each cluster, from negative 100% (all emotions are significantly negatively correlated with BOLD activity in all 5 runs) to positive 100% (all emotions are consistently positively correlated). Thus, inconsistent positive and negative activations across runs cancel each other out. Felt and perceived emotions are shown in blue and red font, respectively. For complete results, see Supplementary Table S7).

## Discussion

Our main finding was that the emotions evoked during free viewing of movies are represented in the brain across numerous spatial scales and patterns. While the self-report data revealed that 46 emotions were consistently used to describe the affective content of the movies, dimension reduction techniques revealed that these could be reduced to eight main emotion dimensions. Neural activity related to both perceived and felt emotions were consistent across individuals and generalized across different stimuli. There was also a region-specific gradient ranging from temporoparietal regions to default mode and subcortical regions for the breadth of emotional tuning. Some brain regions, such as the posterior temporal cortex and cortical midline, were non-selectively activated during most emotions, while the emotion-specific tuning was sharper outside these regions. Even more selective responses to specific emotions were found mainly in subcortical regions. Similarly, there were large differences in the regional specificity of the emotion-evoked activations. While some emotions (such as fear, anxiety, and amusement) consistently engaged large-scale brain networks across temporal, parietal, and midline regions, others, such as anger and disgust, showed more regionally selective patterns. Finally, although perceived and felt emotions were often in alignment, the similarity in emotional perception and feeling and the resultant brain activation varied across emotions. These results constitute one of the most comprehensive investigations of the neural basis of emotional perception and experience. They highlight spatial gradients in the emotion-specificity in the human brain, and outline similarities and differences in the neural basis of perceiving and feeling different emotional states.

### Generalizability and consistency of emotion responses in the brain

The generalizability of emotion models across different samples of the same emotion was highest in temporoparietal, temporal, and anterior cingulate cortices and precuneus. These regions are also activated during the processing of social information ^38^. This generalization confirms that the emotion-dependent responses were not driven purely by visual, auditory, or semantic features of the videos, as these were completely different across the movies. Accordingly, adding low-level stimulus features to the emotion model improved across-stimulus consistency in sensory areas, while the consistency in the regions responding to emotions - temporoparietal, temporal, and midline regions - was unaffected.

The emotion models generalized well across participants, confirming the consistent and shared nature of the emotional responses across individuals also on the neural level. Across-participant generalization of affective responses has been more difficult to achieve than across-stimulus generalization within individuals ^5,6^. However, our results revealed across-participant consistency in widespread brain networks covering the sensory areas (occipital and temporal lobes), posterior midline regions, and parietal regions often associated with attention, consistent with previous studies showing high intersubject synchronization in these regions during emotional moments of movies ^17,38–41^. This likely stems from the high degree of consistency in the time-variant emotional response elicited by the movies, in comparison with static and noisy “snapshot” emotions evoked by e.g. pictures or sound bursts. In movies, emotions occur in natural contexts, which might lead to shared rather than individualistic experiences.

Out of our original 63 emotion categories, 46 (73%) had sufficient occurrence rates and high intersubject reliabilities. The similarity of neural responses revealed that these emotions could be divided into eight clusters. Outside these, we found considerable variability in the reliability and intensity of the emotions evoked by the movies. Emotions such as embarrassment, pride, humiliation, loneliness, shame, gratitude, hurt, and jealousy were difficult to elicit. The common nominator for these emotions is that they often occur in personal social settings, and are not typically experienced in third-person settings, such as while viewing a movie.

### Emotional gradients in the brain

Our results also yielded maps of the affective space in both phenomenological and neural domains. The neural affective space demonstrated a gradient-like organization ranging from generic processing in temporoparietal and default mode regions to more localized, discrete processing in subcortical regions.

The largest and most generic hub of the neural affective space was the temporo-parietal junction (TPJ), which was activated during the perception and experience of most emotions. TPJ is consistently engaged during social perception ^42^ and has been previously shown to contain emotion-specific gradients also during perceived and experienced emotions ^20^. Accordingly, TPJ activation suggests that social information processing is relevant to most emotions evoked by movies. Taken further, social information processing might be an integral part of human emotions in general, which warrants further investigation of the role of social information in emotional processing.

Another central hub for emotion processing was the precuneus, which responded to most perceived and felt emotions. Temporal pole and parietal regions, including SPL and IPL, also responded to numerous emotions. These regions are part of the default mode network that consistently shows differential activity patterns for different emotions ^5,6,18^. The default mode network has been especially associated with sustained, slow changes in emotional states, potentially reflecting the integration of emotion-related information and the conscious experience of emotions ^18,43^.

Some regions showed moderately specific response profiles. For instance, activity in the cerebellum was associated with fear- and surprise-related emotions. Superior parietal regions yielded consistent responses to fear-related emotions. In turn, emotion-evoked activity in subcortical regions was more narrowly tuned and was observed only for some emotions. For example, the amygdala responded primarily to felt and perceived positive emotions and perceived surprise, while the thalamus responded selectively to perceived and felt fear. The selective, local activity does not necessarily mean that these regions would be responsible for a single emotion. Instead, these regions are likely involved in the processing of information that is more pronounced for a subset of emotions. The middle frontal gyrus - implicated in relation to attention orienting - was active during fear-related emotions while showing decreased activity primarily during calm emotional states, potentially reflecting differences in attentional demands between emotions^44^.

Altogether these data show that the emotion circuits in the brain operate in both domain-general and domain-specific manner. The core, domain-general components of the networks likely encode the detailed emotional content of the sensory input, which is then processed in a more granular manner across various extended emotion circuits whose engagement depends on the specific emotional content.

### Cerebral organization of the emotion space

The current study is the first large-scale investigation of emotion clusters identified based on brain activity; the few previous studies have used far smaller sets of emotions ^6^.

The clustering of emotions based on self-reports revealed a primary valence-driven space of emotional feelings, in line with the vast previous literature ^45^. Besides the positively and negatively valenced clusters, we identified four smaller clusters around disgust, annoyance, calmness, and confusion. For 91% of the emotion categories, both perceived and felt emotions from the same category clustered together, most likely reflecting their highly overlapping temporal structure within the movies. The temporal co-occurrence of perception and feeling of the same emotion suggests shared underlying cognitive processes in both.

The two largest and most consistent neural clusters consisted of amusement/confusion- and fear-related emotions, respectively. The amusement/confusion cluster included positive and neutral, mainly arousing emotions. Perceived awkwardness, annoyance, and amusement were clustered together with felt amusement and confusion. The fear-related cluster included various negative emotions, but the most robust activity was detected for fear. Also, four other, smaller clusters (related to joy, romance/neutral, disgust, and calm, positive emotions) showed clear, consistent brain activity. The other clusters (driven mainly by happiness and emotions with a solid bodily component, such as physical pain and sexual desire) showed less consistent activity across stimuli.

The cluster structure based on brain activation was not identical to the structure emerging from the self-report data. In the self-reports, we identified clear emotion clusters (including disgust, fear, and sadness, displeasure, embarrassment, love, joy, and calmness) that resembled those found in previous studies ^16^. Notably, self-reported perceived and felt emotions from the same emotion category often clustered together, while in the neural data, the perceived and felt emotions from the same category often were split into different clusters. As current emotion theories expect ^3,27,46^, subjective experience is only one part of the overall emotional state and does not directly map to the underlying neural state. Thus, emotion-related brain activity reflects various automatic changes in several functional components, including subjective experiences, physiology, motor activation, motivation, and cognition. Our results support the interpretation that neural similarities between emotions result from changes in multiple functional components ^47^.

### Shared and distinct neural basis of perceived and felt emotions

Besides the spatial gradient ranging from generic to specific emotion responses, we also found an anatomical distinction between perceived and felt emotions. Both perceived and felt emotions elicited overlapping brain activity in the precuneus, temporo-parietal junction, and auditory areas. While emotion perception engaged brain regions overlapping with those activated while feeling the same emotion, feelings were accompanied by additional activity in the anterior and prefrontal cortices, thalamus, and parts of the cerebellum, whose activity was absent for emotional perception. These regions are consistently reported to activate during emotional experiences elicited by different types of stimuli ^5,6 36^. Thus, we conclude that the frontal regions, thalamus, and cerebellum process information during felt emotions but not during emotion perception. Especially, we found direct evidence for the involvement of the prefrontal cortex and anterior cingulate in generating subjective experiences of emotions, as also suggested in previous studies ^5^.

Perceived emotion was often accompanied by a concordant feeling, but there were also some emotions for which perception and feeling were decoupled. Alignment between the perception of a character’s emotions and observers’ feelings has been rarely studied. One previous study has shown that characters’ portrayed basic emotions and empathic responses share only 11–35% of the variance in the feelings of the observers ^20^. In our data, we found large variability in the shared variance between felt and perceived emotion from the same categories (average 36%, ranging from almost 0% to close to 100%). Emotion categories related to love and sexual desire, pride and impression, fear, and sadness were both perceived and felt simultaneously. Thus, emotional contagion for these emotions is high, suggesting that the perceived emotion is directly mirrored in the observer. These emotions are characterized by seeking affiliation with others ^48^. On the other hand, the perception of less socially driven emotions such as boredom, interest, confusion, pity, and anger were seldom accompanied by a simultaneous feeling of the same emotion.

### Limitations

We carefully curated a set of 63 emotion categories for the study but could only reliably elicit 46 of them. Especially personal and social emotions, including jealousy, hurt, shame, and gratitude, were difficult to elicit with movies, possibly due to the interpersonal nature of these emotions. Due to methodological constraints (collecting self-reports took 20 hours per participant), we collected self-reports and fMRI data from an independent set of raters. Thus, it is possible that the emotions felt and perceived by the fMRI participants differed from those of the raters. However, the raters came from the same population (young, healthy females) as the fMRI participants and gave consistent ratings. Finally, our emotion models were built on self-reported changes in categorical emotional content, which is optimal for investigating slower changes in conscious perception and feeling but cannot track some of the fast, momentary changes in stimulus content that subcortical regions might be responsible for.

### Conclusions

Our results reveal the neural basis of the core emotional dimensions, and highlight spatial cerebral gradients in representing different emotional states. Eight main dimensions of emotions evoked during movie viewing are encoded in the brain across numerous spatial scales and patterns, and perceived and felt emotions generalize both between individuals and between different stimuli. There is a gradient from large-scale to regionally specific representation of emotions ranging from higher-order temporoparietal areas to default mode network, and finally to more selective activity, especially in subcortical regions. While perception and feeling of different emotions were supported by numerous overlapping brain regions, the activity was more focused in frontal, thalamic, and cerebellar activity during feelings. Although emotional perception and the resultant feelings often go hand in hand, our data highlight that they are also often decoupled in both conscious experience and brain activity.

## Methods

### Generating the high-dimensional emotional space with movies

We initially selected a wide range of emotion categories to cover a high-dimensional affective space (**Figure 1A**). First, we compiled an original list of 128 emotion categories based on previous studies ^6,16,24,49^. Next, we translated the emotion categories into Finnish using a glossary of Finnish emotion words ^50^. We then conducted a pilot study where 25 female volunteers rated the similarity between emotion categories. The ratings were collected using a modified online Q-sort where participants organized the categories into piles based on their felt similarities (https://version.aalto.fi/gitlab/eglerean/sensations; ^51^. Based on the average similarities, we selected a final list of 63 emotion categories that covered the whole emotion space (**Supplementary Table S2**; **Supplementary Figure S1**).

Next, we selected a set of movie stimuli from an existing database of emotional Hollywood movies (**Figure 1B**; ^37^). First, we excluded black-and-white movies to minimize low-level visual differences between scenes. Second, we excluded multiple clips with the same identifiable characters from a single movie. Third, to ensure that the scenes elicit the desired emotions, we collected pilot emotional intensity ratings from 13 Finnish-speaking female volunteers. These raters evaluated the experience of 63 emotion categories (scale: 0=not at all, 4 = extremely much) while viewing the scenes in 3-10-second segments (for more details, see **Supplementary Note 1**). This selection procedure led to a final set of 39 scenes (length 0:16-6:57; total duration 114 minutes; see scene list in **Supplementary Table S3**). For the fMRI study, we divided the movie scenes into five runs of similar emotional content as defined by the pilot ratings (8 scenes in runs 1 –4 and 7 scenes in run 5; for a list of movies in each run and their mean emotion ratings, see **Supplementary Tables S4** and **S5**).

### Generating emotion models from ratings of perceived and felt emotions

We collected dynamic ratings of perceived and felt emotions during movie viewing using an online rating tool from another independent sample of 16 Finnish-speaking female volunteers (**Figure 1C**). The final selection of 39 movie scenes were shown and ratings were collected similarly as in the pilot study. The participants could replay each clip as many times as they wanted. We collected ratings of perceived and felt emotions from the same participants on separate runs and counterbalanced the order of ratings (i.e., perceived or felt first). We created the dynamic emotion models for each movie scene by linear interpolation to the end-points of the short clips. All ratings were set to zero at the beginning of each scene. Due to the low and variable sampling rate and slow changes of the interpolated ratings, we approximated the hemodynamic response function (HRF) as a single gamma function, excluding the undershoot typically included in the canonical double gamma HRFs.

Next, we performed a reliability analysis to ensure that only reliably evoked emotions would be included in the stimulus models (**Figure 1C**). We used a combined reliability measure considering the number of raters that reported perceiving / feeling each emotion, the number of time points when the emotion was detected, and the average intersubject correlation of the emotion ratings. We first calculated the percentage of raters that gave non-zero ratings for each emotion for at least one time point for each run. Second, we calculated the number of time points that received non-zero ratings from at least one rater. Third, we calculated the rating-wise mean intersubject correlation between all pairs of raters. Finally, we calculated the geometric mean of these three measures to derive a joint reliability value for each emotion. To include emotions that were consistently rated for at least one task, we calculated the mean reliability across runs for each task separately and used the higher of these values for each emotion to select the emotions to be included in the subsequent analyses. We evaluated the chance level of reliability as the combined reliability for permuted data. Each permutation consisted of shuffling the values between emotions independently for the three aforementioned measures and recalculating the reliability. The 95th percentile of this distribution across iterations, emotions, and tasks was used as a statistical threshold for defining the reliable emotions used in the data analysis.

### fMRI participants and experimental design

Fifty right-handed Finnish-speaking healthy female volunteers (mean age 24.9±4.4, range 20-38 years) with normal or corrected to normal vision participated in the study. No participants had current psychiatric conditions or medication affecting the central nervous system. All participants gave written informed consent according to the Declaration of Helsinki and were compensated for their time and travel expenses. Aalto University’s ethics committee approved the study protocol.

Functional magnetic resonance imaging (fMRI) was performed on two separate days. In the first fMRI session, we obtained structural images and two functional runs of movie scenes. In the second session, we obtained three functional runs of movie scenes.

Participants watched the movie scenes without a specific task and were instructed only to watch them as they would e.g. watch movies on YouTube. Each of the five runs consisted of 7-8 movie scenes and lasted for approximately 23 minutes (range 22-24 mins). The order of movie scenes within a run was fixed for all participants, and the order of runs was counterbalanced across participants. A run started with a fixation cross presented for 9.6 seconds (i.e., 4 TRs), followed by the first scene. The movies were played one after another without gaps. After the last movie of a run, a fixation cross was presented for 9.6 seconds.

Sound was delivered through Sensimetrics S14 insert earphones (Sensimetrics Corporation, Malden, Massachusetts, USA). Each participant’s sound level was adjusted individually to be loud enough over the scanner noise. Visual stimuli were back-projected on a semi-transparent screen using a Panasonic PT-DZ110XEJ data projector (Panasonic, Osaka, Japan) and via a mirror to the participant. Stimulus presentation was controlled with Presentation software (Neurobehavioral Systems Inc., Albany, CA, USA). An fMRI-compatible face camera (MCR Systems Ltd, Leicester, UK) was used to ensure that participants had their eyes open during the whole scan.

### fMRI data acquisition and preprocessing

We collected MRI data with a 3T Siemens Magnetom Skyra scanner at the Advanced Magnetic Imaging Centre, Aalto Neuroimaging, Aalto University. To improve the binocular field of view, we used a 30-channel Siemens receiving head coil modified from a standard 32-channel coil by removing the two coil elements surrounding the eyes. We collected whole-brain functional scans using a whole-brain T2*-weighted EPI sequence with the following parameters: 44 axial slices, interleaved order (odd slices first), TR = 2.4 s, TE = 24 ms, flip angle = 70°, voxel size = 3.0 × 3.0 × 3.0 mm^3^, matrix size = 64 × 64 corresponding to FOV 192 × 192 mm^2^, PAT2 parallel imaging. To avoid the signal from fat tissue, we used a custom-modified bipolar water excitation radio frequency (RF) pulse. Finally, we collected high-resolution anatomical images with isotropic 1 × 1 × 1 mm^3^ voxel size using a T1-weighted MP-RAGE sequence.

For each BOLD-fMRI run, we performed the preprocessing with *fMRIprep* version 1.1.8 (^52^; RRID:SCR_016216). First, we used *fMRIprep* to prepare a reference volume and its skull-stripped version. Next, we co-registered the BOLD reference to the T1w reference image using *bbregister* (FreeSurfer), which implements boundary-based registration ^53^. Co-registration was configured with nine degrees of freedom to account for distortions remaining in the BOLD reference. Head-motion parameters for the BOLD reference (transformation matrices and six rotation and translation parameters) were estimated before spatiotemporal filtering using *mcflirt* (FSL 5.0.9, ^54^). BOLD runs were slice-time corrected using *3dTshift* from AFNI (^55^, RRID:SCR_005927). The resulting slice-timing corrected BOLD time series were resampled onto their original, native space by applying a single, composite transform to correct head-motion and susceptibility distortions. Next, we applied spatial smoothing with an isotropic, Gaussian kernel of 6mm FWHM (full-width half-maximum). The resulting “non-aggressively” denoised runs were produced, and the noise regressors were used for nuisance regression during analysis. The BOLD time series were resampled to MNI152NLin2009cAsym standard space, generating a preprocessed BOLD run in the corresponding space.

After preprocessing, we extracted the voxel-wise BOLD time series within the group brain mask, and converted them to percent signal change for each participant. Next, we calculated the mean ROI time series in regions defined with Brainnetome ^56^ and cerebellar connectivity ^57^ atlases covering the cerebral cortex, subcortical regions, and the cerebellum. We included 273 regions in the analyses after omitting one cerebellar ROI falling outside the group brain mask. Signals extracted from deep white matter and cerebrospinal fluid and the six motion time series and their second-order effects and linear trends were regressed from the ROI level data. To match the frequency content of the slowly evolving emotion time series (emotion event lengths ranged from 20 to 100 seconds), we filtered the data of periods between 200 (twice the length of the longest event) and 20 seconds (i.e., .005 to .05 Hz). We employed a finite impulse response filter designed with the Parks-McLellan algorithm with a ripple of <1dB in the passband and 40dB in the stopband. Transition bands extended from ∼–24dB at 0Hz in the lower stopband and up to 0.0613Hz in the upper stopband.

### Fitting the emotion models

We estimated the beta weights of the emotion models with least-squares fitting after removing the mean of the signals and models (**Figure 1D**). For between-runs generalization, we fitted the model to the mean standardized activity of all participants in the training runs (three runs). We then tested the model fit on the individual activity time series of the participants in the test runs (the remaining two runs). The process was repeated for all permutations of training and test runs (10 combinations in total). Next, for within-run generalization, we used 10-fold cross-validation across participants. Here, we fitted the model to the mean activity of 45 participants, and the fit of the predicted activity was tested on the individual activity of the left-out participants. This process was repeated for each of the ten cross-validation folds and ten combinations of runs.

To test whether low-level stimulus features could predict the fit, we repeated the same fitting process for the emotion models extended with two visual features (presence of high spatial frequencies and differential energy between subsequent frames) and one auditory feature (root-mean-squared power) from the movies. The resulting low-level stimulus feature time series were added to a combined emotion and low-level stimulus features model.

### Reliability of the neural responses to emotion categories

To evaluate the reliability of emotion responses while avoiding the circularity of correlating neural responses within the same run, we calculated a spatial correlation matrix for each emotion between all combinations of non-overlapping training and test runs used in the between-runs cross-validation (i.e. emotion responses in the training set were correlated with those in the test set). A mean correlation matrix was produced by averaging the correlation matrices over the 10 run combinations, and this matrix was used for evaluating the reliability of brain-wide responses to emotions and performing the subsequent brain-based clustering. Additionally, to reduce the effects of non-significant activations on the brain-wide reliability measure, we repeated a similar analysis using generalized (two-sided) Dice indices of the thresholded activity maps. This procedure provided highly similar results as the correlation matrices and is therefore omitted in the subsequent sections.

### Cluster analysis

To investigate the similarity structure of the dynamic emotion ratings and emotion-related brain activity, we performed an average linkage hierarchical clustering for both data separately. The leaf order was optimized by maximizing the sum of the similarities of adjacent emotions (optimalleaforder-function, https://www.mathworks.com/help/stats/optimalleaforder.html), and default thresholding was used (70% of maximum linkage) as the clustering cutoff for the visualization.

## Acknowledgments

We thank Marita Kattelus for her help with the data acquisition. This work was supported by the Academy of Finland (grants 323425 to HS and 276643 and 332398 to IPJ) and the Finnish Cultural Foundation (grant 150496 to JML). This research is also part of the project Right to Belong which is funded by the Strategic Research Council (352648 and 352655).

## Contributions

Conceptualization and writing by all authors; Funding acquisition by IPJ; Supervision by JML, LN, IPJ, and MS; Data acquisition by SV, HS, and AA; Methodology, data analyses, and visualization by JML, HS, SV, and SS.

