## Supplementary Information for "Cerebral Topographies of Perceived and Felt Emotions"

### Supplementary Note 1

To enable parallel near-continuous rating of all emotion categories, each movie scene was cut into 3-10-seconds long clips based on the original cut-points preferring local loudness minima in the soundtrack of the movies to avoid splitting the videos in the middle of continuous actions. Where the time between subsequent cuts exceeded the maximum clip duration of 10 seconds, the cut interval was first equally split into the minimum number of <10-second windows. The splitting points were then adjusted to coincide with the closest loudness minima of the soundtrack. The raters evaluated the intensity of each emotion elicited by the clip using a Likert scale from 0 to 4. Finally, we calculated the average intensity of each emotion category for each movie scene by averaging across the scene-wise clips, arranged movie scenes by the most intense emotion categories they elicited, and manually selected the scenes to cover a wide and equal range of different emotion categories (see Supplementary Tables S1 and S2).

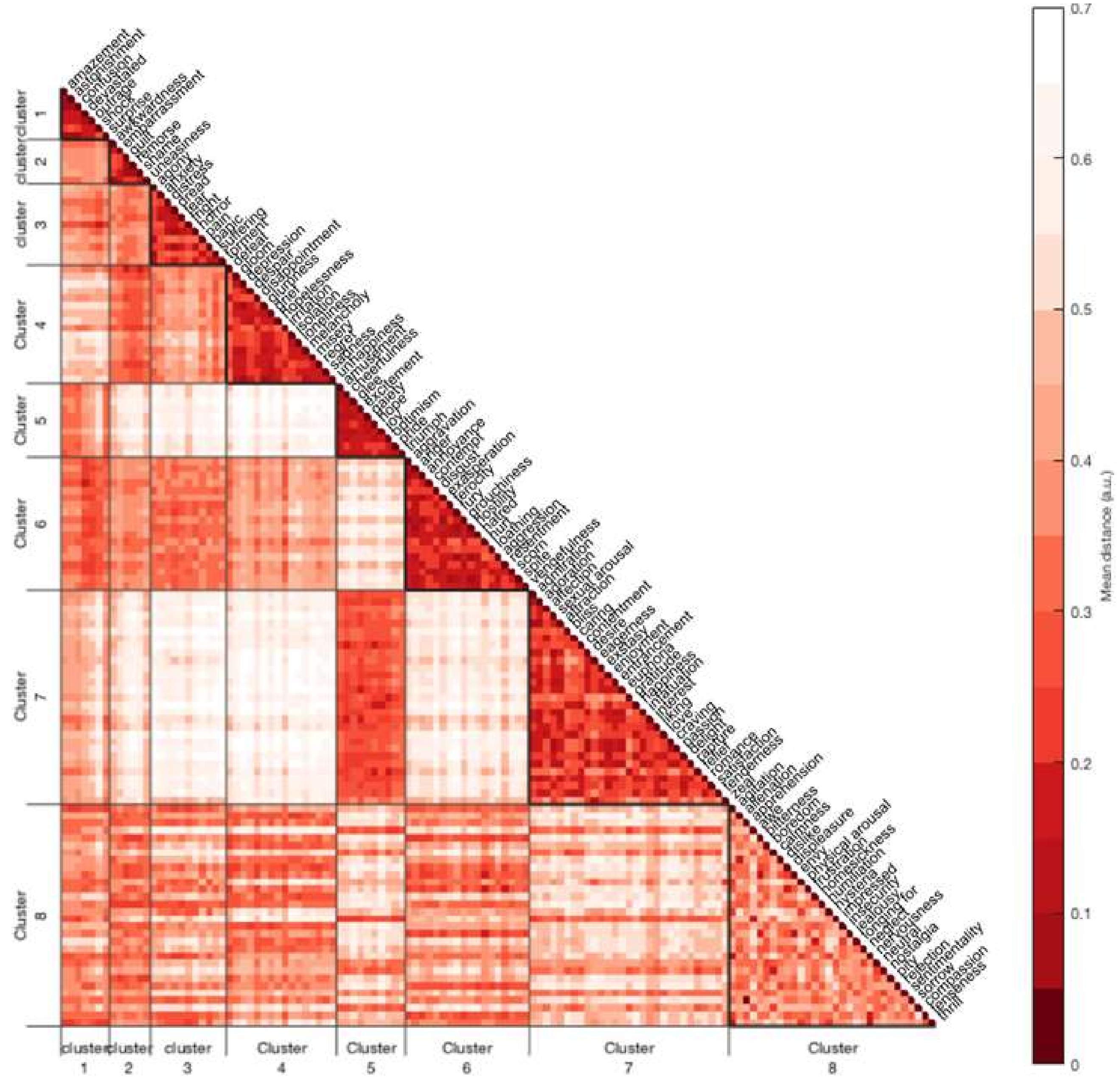

**Supplementary Figure S1.**

**The similarity of emotion categories.** To aid the selection of emotion categories that span the entire emotion space, we ran a pilot study (N=25 participants) to collect similarity ratings between our original list of 127 emotion categories. The ratings were collected using an online sorting tool (https://version.aalto.fi/gitlab/eglerean/sensations; initially used in Nummenmaa, L., Hari, R., Hietanen, J. K., & Glerean, E. (2018). Maps of subjective feelings. *Proceedings of the National Academy of Sciences*, *115*(37), 9198-9203). Clusters were identified with density-based clustering (DBSCAN; Ester, M., Kriegel, H. P., Sander, J., & Xu, X. (1996) A density-based algorithm for discovering clusters in large spatial databases with noise. In *kdd* (Vol. 96, No. 34, pp. 226-231)).

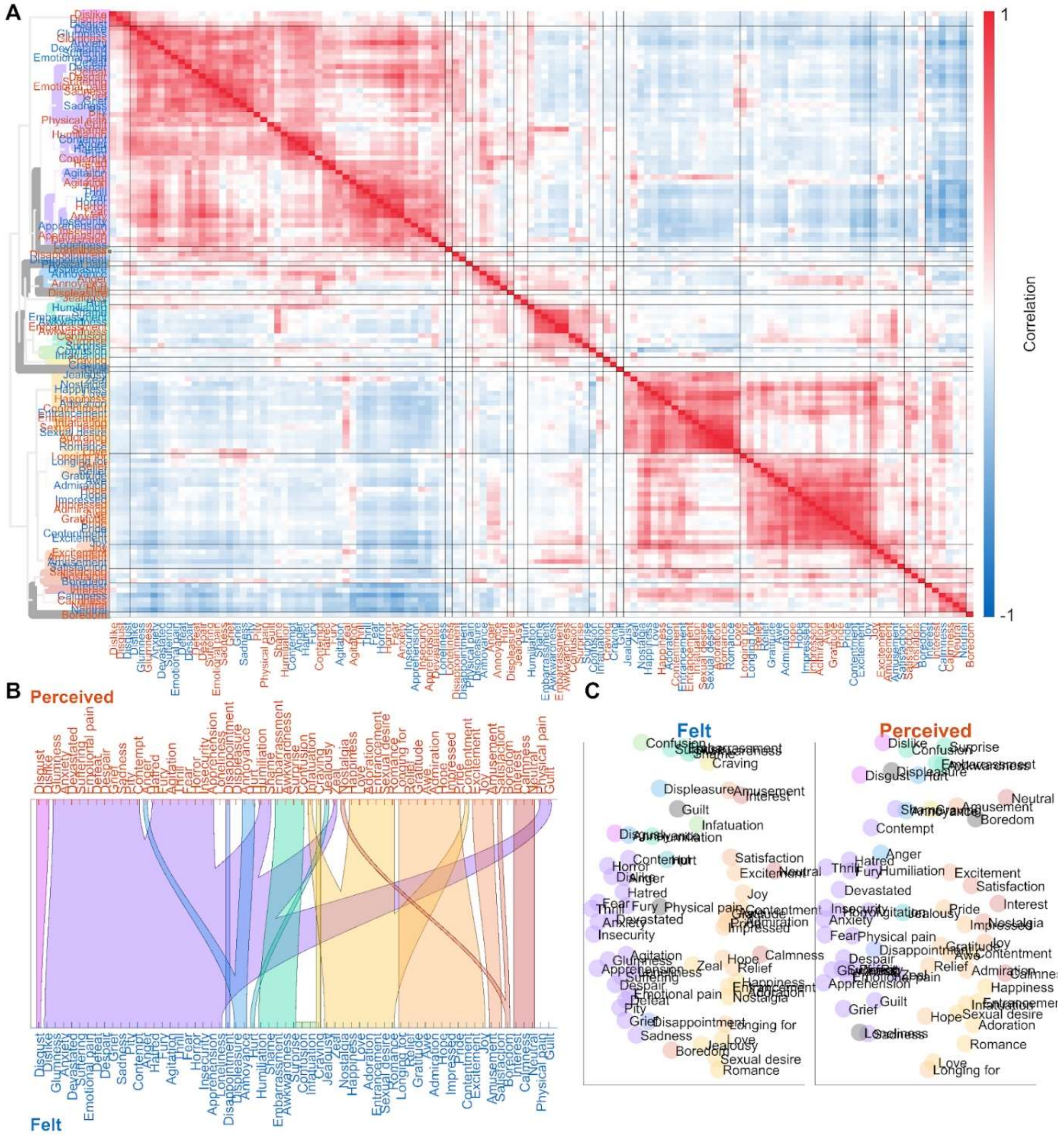

**Supplementary Figure S2.**

**Cluster structure for all emotions based on ratings. A** Correlation matrix and dendrogram of ratings over emotion models across all runs and both felt and perceived emotions in the set of reliable emotion ratings. **B** Alluvial diagram shows cluster labels' correspondence between felt and perceived emotions. **C** Multidimensional scaling visualizes the similarity of emotions based on only the ratings of felt emotion (top) and perceived emotions (bottom). The emotions that did not belong to any cluster are shown in gray in the dendrogram and multidimensional scaling plots and are left unconnected in the alluvial diagram.

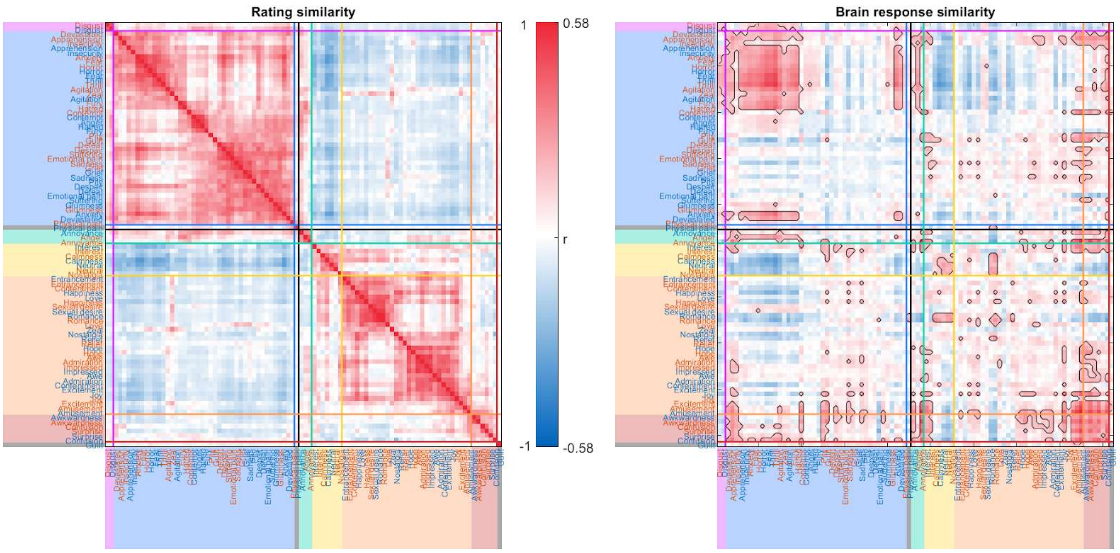

### Supplementary Figure S3.

**Similarity between emotion categories.** The similarity between emotion categories is shown for emotion ratings (left) and cross-validated reliability and spatial correlation of brain responses to the emotions (right). Emotions are organized by the rating clusters in Figure 2. Significant similarity of brain responses compared with randomly sampled data (p<.05, uncorrected) is indicated by black contours.

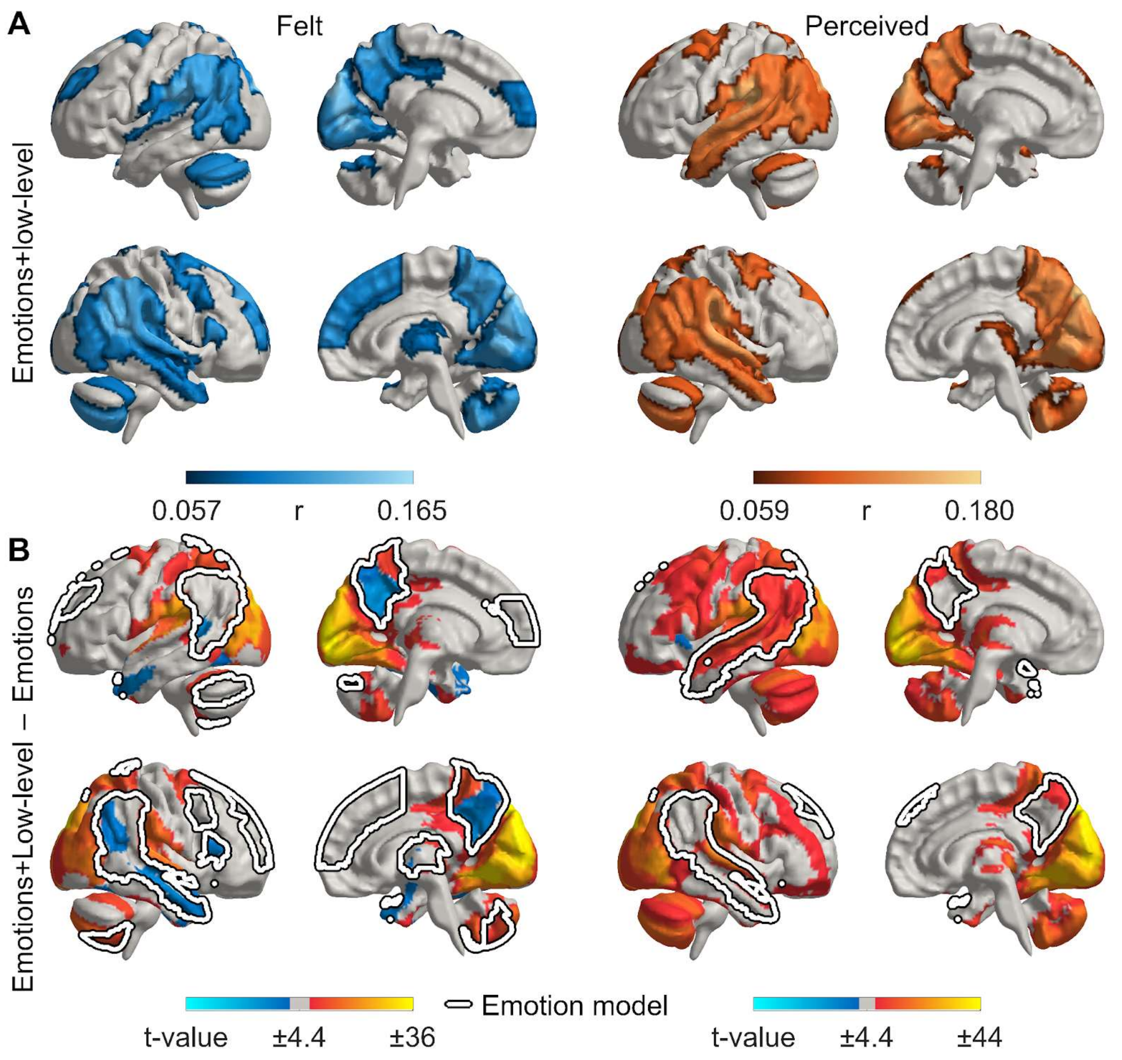

**Supplementary Figure S4.**

**Generalization of emotion models extended with low-level stimulus features across runs. A** Model fits for the extended emotion model. Results are analogous to Fig. 3B, but in addition to emotion models, low-level visual (amount of high spatial frequencies and differential energy between subsequent frames) and auditory (root-mean-squared power) features were added into the model. **B** Contrast of model fits for the extended emotion+low-level models vs. emotion-only models. Hot colors indicate where the low-level stimulus features improved model fit, cold colors indicate areas where stimulus features decreased the cross-validated model fit, presumably due to overfitting the training data. The contrast is thresholded at $p<.05$, Bonferroni corrected over ROIs and the two emotion models (corrected $p=.05/273/2$). The outlines indicate areas where the between-runs model fit was significant for the emotion models in Fig. 3B.

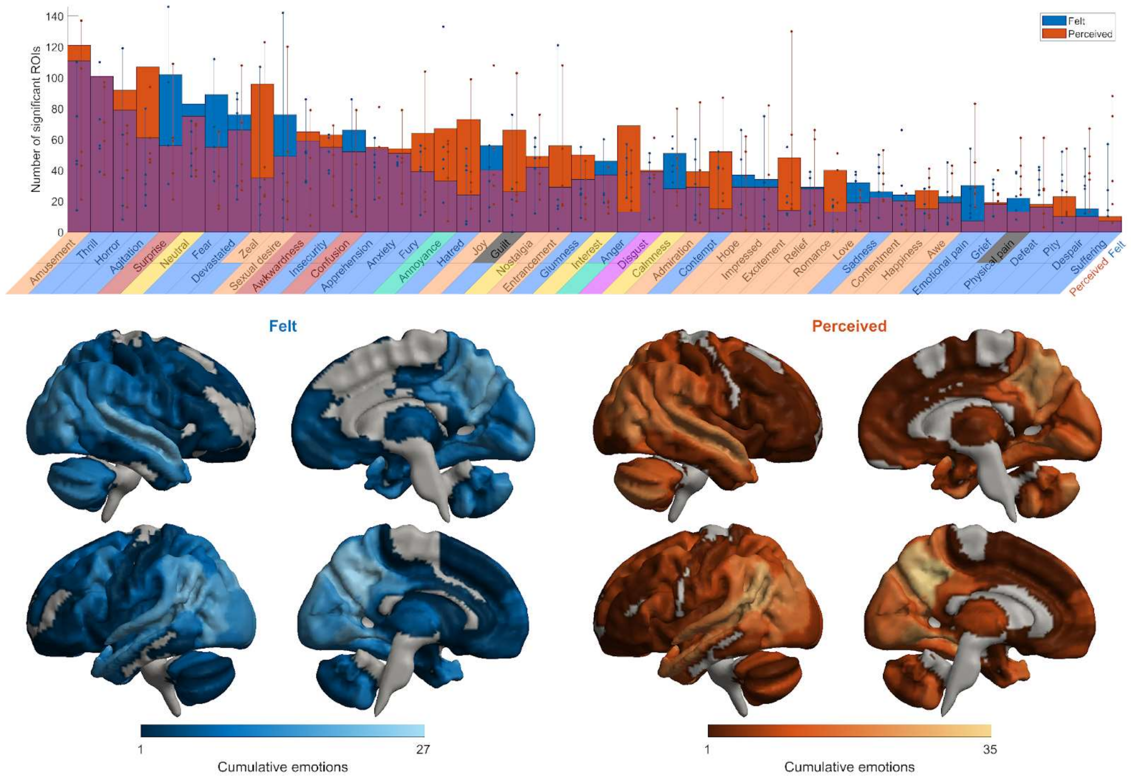

**Supplementary Figure S5.**

**Total brain area activated by individual emotions. Top:** Total brain area activated by individual emotions (**top**) colored by clusters derived from spatial clustering of neural responses. **Bottom**: Cumulative maps show the number of emotions whose responses were statistically significant in at least one run. The data are thresholded at two-tailed $p < .05$ (FWER corrected).

**Supplementary Table S1.**

Region-wise correlations for the cross-validated emotion model fitting.

| | Centroid MNI coordinates (mm) | | | | |
|---|---|---|---|---|---|
| ROI | X | Y | Z | Felt corr. (mean+-std) | Perceived |
| SFG_R | 9.5 | 19.3 | 57.2 | 0.07+-0.05 | |
| SFG_L | -8.6 | 52.3 | 42.4 | | 0.06+-0.05 |
| SFG_R | 15.7 | 51.5 | 43.1 | 0.10+-0.04 | 0.07+-0.04 |
| SFG_R | 8.8 | 41.5 | 38.1 | 0.06+-0.05 | |
| SFG_L | -4.6 | 59.6 | 18.0 | 0.06+-0.04 | |
| SFG_R | 10.6 | 61.7 | 15.7 | 0.08+-0.05 | |
| MFG_L | -25.0 | 46.0 | 33.8 | 0.06+-0.04 | |
| MFG_R | 45.2 | 14.5 | 41.5 | 0.06+-0.05 | |
| MFG_R | 30.7 | 58.6 | 19.6 | 0.07+-0.04 | |
| IFG_R | 57.2 | 26.9 | 14.0 | 0.09+-0.05 | |
| STG_L | -60.0 | -30.0 | 9.6 | | 0.08+-0.02 |
| STG_R | 69.4 | -17.1 | 8.7 | | 0.06+-0.02 |
| STG_R | 50.1 | 15.6 | -17.3 | 0.07+-0.03 | 0.07+-0.04 |
| STG_L | -52.5 | -0.2 | -7.9 | | 0.09+-0.03 |
| STG_R | 59.4 | -9.0 | -2.5 | | 0.08+-0.03 |
| MTG_L | -51.6 | 4.4 | -25.3 | | 0.08+-0.04 |

| | | | | | |
|---|---|---|---|---|---|
| MTG_R | 55.1 | 8.2 | -28.2 | 0.07+-0.04 | 0.09+-0.05 |
| MTG_L | -56.7 | -54.9 | 7.0 | 0.08+-0.04 | 0.07+-0.03 |
| MTG_R | 63.3 | -50.9 | 5.3 | 0.07+-0.05 | |
| MTG_L | -56.1 | -16.6 | -7.1 | | 0.09+-0.04 |
| MTG_R | 61.1 | -13.1 | -7.4 | 0.07+-0.03 | 0.08+-0.03 |
| pSTS_L | -52.2 | -37.2 | 6.7 | | 0.09+-0.04 |
| pSTS_R | 55.7 | -34.2 | 6.1 | 0.08+-0.03 | 0.09+-0.04 |
| pSTS_L | -50.1 | -47.5 | 13.4 | 0.08+-0.03 | 0.07+-0.03 |
| pSTS_R | 59.9 | -37.4 | 15.4 | 0.11+-0.03 | 0.06+-0.03 |
| IPL_L | -53.5 | -46.7 | 40.2 | 0.10+-0.04 | 0.07+-0.04 |
| IPL_R | 60.3 | -40.9 | 41.0 | 0.11+-0.05 | 0.07+-0.04 |
| IPL_L | -44.2 | -62.5 | 28.8 | 0.08+-0.04 | 0.07+-0.03 |
| IPL_R | 56.1 | -51.8 | 27.5 | 0.12+-0.04 | 0.08+-0.03 |
| Pcun_L | -2.3 | -61.0 | 54.0 | 0.10+-0.04 | 0.10+-0.04 |
| Pcun_R | 8.9 | -62.5 | 53.9 | 0.10+-0.04 | 0.09+-0.03 |
| Pcun_L | -5.1 | -44.7 | 60.1 | 0.07+-0.03 | |
| Pcun_R | 10.3 | -44.5 | 61.5 | 0.06+-0.03 | |
| Pcun_L | -3.3 | -52.1 | 36.5 | 0.08+-0.04 | 0.09+-0.04 |
| Pcun_R | 9.3 | -51.3 | 37.5 | 0.09+-0.03 | 0.10+-0.04 |
| Tha_R | 9.5 | -8.5 | 8.4 | 0.06+-0.04 | |

| | | | | |
|---|---|---|---|---|
| Tha_R | 12.4 | -11.3 | 16.9 | 0.06+-0.04 |
| Cer_L | -33.7 | -66.6 | -29.7 | 0.07+-0.04 |
| Cer_L | -23.8 | -73.6 | -39.8 | 0.08+-0.03 |
| Cer_V | 1.2 | -66.0 | -29.1 | 0.06+-0.03 |
| Cer_V | 2.3 | -65.1 | -35.9 | 0.06+-0.04 |
| Cer_V | 1.9 | -61.4 | -39.5 | 0.06+-0.04 |

**Supplementary Table S2.**

List of emotion categories. Green = emotion categories chosen based on earlier studies ([1] Cowen & Keltner 2017; [2] Skerry & Saxe 2015; [3] Saarimäki et al. 2018; [4] Adolphs 2002). Yellow = emotion categories added based on pilot similarity ratings (N=25) from the emotion list by Shaver et al. (1987).

| Cluster | Emotion word (FIN) | Emotion word (ENG) | Primary reference |
|---|---|---|---|
| 1 | hämmennys | confusion | 1 |
| 1 | pöyristyneisyys | devastated | 2 |
| 1 | hämmästys | surprise | 1-5 |
| 2 | kiusaantuneisuus | awkwardness | 1 |
| 2 | nolous | embarrassment | 2,4-5 |
| 2 | syyllisyys | guilt | 2-5 |
| 2 | häpeä | shame | 2.5 |
| 3 | ahdistus | anxiety | 1.5 |
| 3 | pelko | fear | 1, 3-5 |
| 3 | kauhu | horror | 1, 5 |
| 3 | fyysinen kipu | physical pain | 1 |
| 3 | henkinen kipu | emotional pain | 1 |
| 3 | kärsimys | suffering | 5 |
| 4 | häviäminen | defeat | 5 |
| 4 | epätoivo | despair | 3, 5 |
| 4 | pettymys | disappointment | 2, 5 |
| 4 | synkkyys | glumness | 5 |
| 4 | murhe | grief | 5 |
| 4 | yksinäisyys | loneliness | 2, 5 |
| 4 | suru | sadness | 1, 3-5 |
| 5 | huvittuneisuus | amusement | 1, 5 |
| 5 | innostus | excitement | 1-2, 5 |
| 5 | toiveikkuus | hope | 2, 5 |
| 5 | ilo | joy | 1-2, 5 |
| 5 | ylpeys | pride | 2-5 |
| 6 | suuttumus | anger | 1, 3-5 |
| 6 | ärtymys | annoyance | 2, 5 |
| 6 | paheksunta | contempt | 3-5 |
| 6 | inho | disgust | 1-5 |
| 6 | raivo | fury | 2, 5 |
| 6 | viha | hostility | 5 |
| 6 | loukkaantuneisuus | hurt | 5 |
| 7 | ihailu | admiration | 1, 4 |
| 7 | ihastus | adoration | 1, 5 |
| 7 | tyytyväisyys | contentment | 2, 5 |
| 7 | halu | sexual desire | 1 |
| 7 | lumoutuneisuus | entrancement | 1 |
| 7 | kiitollisuus | gratitude | 2.3 |
| 7 | onnellisuus | happiness | 3-5 |
| 7 | hullaantuminen | infatuation | 4.5 |
| 7 | kiinnostus | interest | 1 |

| Cluster | Emotion word (FIN) | Emotion word (ENG) | Primary reference |
|---|---|---|---|
| 7 | rakkaus | love | 2,4-5 |
| 7 | himo | craving | 1 |
| 7 | helpotus | relief | 1, 5 |
| 7 | romanttisuus | romance | 1 |
| 7 | tyydytys | satisfaction | 1 |
| 7 | kiihko | zeal | 5 |
| 8 | huolestuneisuus | apprehension | 2, 5 |
| 8 | syvä kunnioitus | awe | 1 |
| 8 | ikävystyminen | boredom | 1 |
| 8 | rauhallisuus | calmness | 1 |
| 8 | vieroksuminen | dislike | 5 |
| 8 | tyytymättömyys | displeasure | 5 |
| 8 | kiihtymys | emotional arousal | 5 |
| 8 | nöyryytys | humiliation | 5 |
| 8 | vaikuttuneisuus | impressed | 2 |
| 8 | epävarmuus | insecurity | 5 |
| 8 | mustasukkaisuus | jealousy | 2, 5 |
| 8 | kaipaus | longing for | 3, 5 |
| 8 | neutraali | neutral | 3 |
| 8 | nostalgia | nostalgia | 1, 2 |
| 8 | sääli | pity | 5 |
| 8 | jännitys | thrill | 5 |

**Supplementary Table S3.**

Content and duration of the movie clips.

| Clip number | Original movie | Duration | Content |
|---|---|---|---|
| 3 | The dead poet's society | 4:14 | Todd (Ethan Hawke) commits suicide. |
| 7 | Seven | 1:43 | Crime linked to the sin of sloth. A man apparently dead is discovered lying on a bed, his hands tied. He is extremely skinny, and has been savagely tortured. The word “Sloth” has been written on the walls, and in the room they find pictures depicting different stages of the victim’s progressive demise. Unexpectedly, the man wakes up. |
| 9 | E.T. | 4:35 | E.T. is going to die, surrounded by scientists. |
| 10 | Trainspotting | 1:37 | In an apartment, several persons are sleeping. Then, a women screams. “Sick Boy” (played by Johnny Lee Miller) tries to calm her down. In the meantime, the others wake up. They eventually find out that the woman’s newborn baby is dead. After a long silence, Sick boy asks Mark (Ewan McGregor) to say something. Mark then says he will make a “fix”. |
| 14 | When Harry met Sally | 2:45 | In a very well-known scene, Sally (Meg Ryan) fakes an orgasm in the restaurant, provoking Harry’s (Billy Cristal) embarrassment. |
| 15 | Forrest Gump | 2:01 | The child is introduced to Forrest. The boy goes to sit down in front of the television. Jenny – the mother – tells Forrest that this is his son. Forrest sits down close to the boy. Very moved, Jenny watches the father and the son sitting close to each other. |
| 16 | Scream | 6:33 | A girl answers to the phone. She is asked what her preferred horror movie is. Progressively, she finds out that the person she is speaking to knows her, and has a serious intention to kill her. Afterwards, she sees her boyfriend being killed by this person. Eventually, the killer, wearing a black coat and a grotesque mask, gets into the house and chases her with a knife. At the end, he manages to kill her. |
| 17 | Trainspotting | 1:02 | After a drunken night, Spud (Ewen Bremmer) wakes up in his girlfriend’s bed, and realizes that the sheets are dirty with his excrements. In the following scene, he tries to hide this from the gilfriend’s mother, who wants to take the sheets for the laundry. In the confusion, the mother pulls the sheets away from Spud’s hands, and accidentally splashes the whole family with excrements. |
| 18 | The professional | 2:44 | Léon (Jean Reno) plans the escape of Mathilda (Nathalie Portman). He puts her in the ventilation circuit. She doesn’t want to leave Leon. He promises that they are going to reunite later. They say goodbye to each other, and Mathilda understands that she is never going to see Leon again. |
| 21 | Dead man walking | 6:40 | Execution of Matthew by injection (Sean Penn): He is tied on the execution table, and the scene shows the lethal substance being progressively injected in his veins. |
| 22 | The silence of the lambs | 3:29 | Extraction of a butterfly’s larva from a dead body’s mouth. |
| 23 | A fish called Wanda | 2:53 | Archie (John Cleese) gets undressed, waiting for his girlfriend. Unexpectedly, the owners of the house get into the house and discover him naked. |
| 25 | Sleepers | 2:20 | The guardian brings the children to the cave, so that he can sexually abuse them. |
| 26 | When a man loves a woman | 1:39 | Alice (Meg Ryan) promises Michael (Andy Garcia) to never acting impulsively again, and she promises to stop drinking. |
| 27 | Saving private Ryan | 5:22 | Beginning of the movie. In Omaha Beach, American troops landing in June 1944. A heavy fighting unfolds, where several soldiers are killed amid several horrible scenes. |
| 28 | The shining | 4:15 | Jack (Jack Nicholson) pursues his wife with an axe. |
| 30 | In the name of the father | 3:30 | Violent interrogation of Gerry (Daniel Day-Lewis), when the interrogators threaten to murder his father. He eventually signs a confession forged by the interrogators. |
| 31 | Indiana Jones and the last crusade | 1:53 | Indiana Jones (Harrison Ford) escapes from a catacomb full of rats, in Venice, under a library. |
| 32 | Copycat | 2:23 | Monahan (Holly Hunter) goes to the toilets to looks for the murderer; she gets caught. |

| Clip number | Original movie | Duration | Content |
|---|---|---|---|
| 33 | Ghost | 3:35 | Molly (Demi Moore) and Sam (Patrick Swayze) make pottery together, in a very romantic scene. |
| 35 | Trainspotting | 1:44 | Mark (Ewan McGregor) is a drug addict who has not taken heroin from a while and is suffering from withdrawal symptoms. As a consequence, he suffers from a violent diarrhoea. He is then obliged to go to an extremely dirty public restroom. After defecating, he remembers that he had just hidden a newly purchased heroin pill in his anus. He is then forced to search deep through his excrements for the pill. |
| 36 | City of Angels | 4:15 | Maggie (Meg Ryan) dies in Seth's (Nicolas Cage) arms. |
| 38 | IT | 2:13 | A clown hidden in the sewer attracts a boy. |
| 39 | The piano | 0:43 | Stewart (Sam Neil) cuts off Ada's hand (Holly Hunter) with an axe. |
| 43 | A perfect world | 4:27 | Butch (Kevin Costner) is gunned down, at the end of the movie. |
| 46 | Child's play II | 1:05 | Chucky beats Andy's teacher with a ruler. |
| 48 | Life is beautiful | 3:48 | In a prisoner's camp, the father (Roberto Benigni) translates the orders given by the soldier to the prisoners. He is not actually translating, but he is making up a translation that does not scare his son. Specifically, he is trying to make his son believe that all this is a large-scale game. |
| 49 | Blue | 0:25 | A woman goes up on an escalator, carrying a box. |
| 50 | Misery | 3:31 | Annie (Kathy Bates) breaks Paul's legs (James Caan). |
| 51 | Leaving Las Vegas | 2:30 | Sera (Elisabeth Shue) is raped and beaten by 3 young men. |
| 52 | Dangerous minds | 2:08 | The character played by Michelle Pfeiffer tells the class that one of their classmates is dead. |
| 53 | Underground | 0:58 | A man's attempts to have sex with a prostitute are interrupted by the bombing of the city by German planes during WWII. |
| 55 | The Blair witch project | 3:57 | Anxious-provoking scene by the end of the movie: Heather (Heather Donahue), and Mike (Michael Williams) – who is filming Heather - are looking for Joshua (Joshua Leonard) in the woods, at night. They hear screams, apparently form Joshua. They find a house, from where the screams are coming. Mike and Heather go to the second floor. Next, Mike comes back without Heather. Heather's screams can be heard. Eventually, Mike's camera falls on the ground, and keeps filming. |
| 56 | Benny & Joone | 2:01 | Benny (Johnny Depp) plays the fool in a coffee shop. |
| 57 | Hellraiser | 1:30 | On the floor, the size of two stains are growing, and progressively transforming into a monster with a human-like skeleton. |
| 58 | The lover | 0:43 | Marguerite (Jane March) gets into a car, and the car starts to ride. She is dropped of on an animated street. She knocks on a door, and a Chinese man opens and lets her in. |
| 59 | The dead poet's society | 2:40 | By the end of the movie, all the students climb on their desks to manifest their solidarity with Mr. Keating (Robin William), who has just been fired. |
| 61 | There is something about Mary | 2:55 | Ted (Ben Stiller) fights with the dog. |
| 62 | Philadelphia | 5:28 | Andrew (Tom Hanks) and Joe (Denzel Washington) listen to an opera aria on the stereo. Ted describes to Joe the pain and passion felt by the opera character. |
| 64 | Blue | 0:16 | A person passes a piece of aluminium foil through the window of a car. |

Due to their large size, the following supplementary materials are shared online with a permanent identifier:

**Supplementary Table S4.**

Mean felt emotion intensity ratings (range 0-4) for each movie scene. For a downloadable, color-annotated table, see 10.6084/m9.figshare.20418489.

**Supplementary Table S5.**

Mean perceived emotion intensity ratings (range 0-4) for each movie scene. For a downloadable, color-annotated table, see 10.6084/m9.figshare.20418489.

**Supplementary Table S6.**

Emotion-specific consistency across runs for each region of interest. For a downloadable table, see 10.6084/m9.figshare.20418489.

**Supplementary Table S7.**

Mean *t* values for each region of interest ordered by cluster ($p<.05$, Bonferroni corrected over regions and emotions). For a downloadable table, see 10.6084/m9.figshare.20418489.
